# Evaluating computational tools for protein-coding sequence detection: Are they up to the task?

**DOI:** 10.1101/2024.05.16.594598

**Authors:** DJ Champion, Ting-Hsuan Chen, Susan Thomson, Michael A. Black, Paul P. Gardner

## Abstract

**Background:** Detecting protein-coding genes in nucleotide sequences is a significant challenge for understanding genome and transcriptome function, yet the reliability of bioinformatic tools for this task remains largely unverified. This is despite some tools being available for several decades, and widely used for genome and transcriptome annotation.

**Results:** We perform an assessment of nucleotide sequence and alignment-based *de novo* protein-coding detection tools. The controls we use exclude any previous training dataset and include coding exons as a positive set and length-matched intergenic and shuffled sequences as negative sets.

Our work demonstrates that several widely used tools are neither accurate nor computationally efficient for the protein-coding sequence detection problem. In fact, just three of nine tools significantly outperformed a naive scoring scheme. Furthermore, we note a high discrepancy between self-reported accuracies and the accuracy achieved in our study. Our results show that the extra dimension from conserved and variable nucleotides in alignments have a significant advantage over single sequence approaches.

**Conclusions:** These results highlight significant limitations in existing protein-coding annotation tools that are widely used for lncRNA annotation. This shows a need for more robust and efficient approaches to training and assessing the performance of tools for identifying protein-coding sequences. Our study paves the way for future advancements in comparative genomic approaches, and we hope will popularise more robust approaches to genome and transcriptome annotation.

## Introduction

Annotating protein-coding regions within genomes and transcripts remains a crucial bioinformatic challenge [Fickett and Tung, 1992, Amaral et al., 2023, Zerbino et al., 2020], particularly in the context of differentiating between protein-coding and long non-coding RNA transcripts [Xu et al., 2017, Li et al., 2020]. However, it remains unclear how well the available tools handle realistic datasets, including truncations, sequencing, and processing errors [Zerbino et al., 2020].

In model species like *Homo sapiens, Mus musculus, Drosophila melanogaster*, and *Saccharomyces cerevisiae*, rigorous manual biocuration has significantly refined genome annotations [Frankish et al., 2023, Larkin et al., 2021, Engel et al., 2022]. For example, *H. sapiens* genome annotations have evolved from an initial estimate of 30,000 protein-coding genes to about 20,000 through decades of meticulous review, which highlights the importance of cautious interpretation of predicted annotations [Hatje et al., 2019]. These annotations have been enhanced by experimental methods that detect active gene expression in specific tissues and conditions, though such methods are limited by experimental parameters and can include noisy non-functional transcription and translation signals [Hatje et al., 2019, Zerbino et al., 2020, Palazzo and Lee, 2015, Mudge et al., 2022, Alexander F. Palazzo and Kang, 2024].

As genome sequencing becomes more accessible, manual annotation, the gold standard, has been increasingly replaced by automated techniques for genome and transcriptome annotation[Guigó et al., 2006, Zerbino et al., 2020, Stein, 2001, Tang et al., 2019]. Despite extensive literature on genome annotation [Pinkney et al., 2020, Li et al., 2020, Klapproth et al., 2021, Dindhoria et al., 2022, Amaral et al., 2023], there are few systematic evaluations of tools that differentiate coding from non-coding sequences [Fickett and Tung, 1992, Guigó et al., 2006, Klapproth et al., 2021, Zheng et al., 2021, Singh and Roy, 2022]. Therefore, we believe that it is timely for further evaluations of genome annotation tools.

The field of genome annotation could greatly benefit from unbiased benchmarking protocols similar to those used in protein structure prediction, such as those in the Critical Assessment of Protein Structure Prediction (CASP) initiative [Kryshtafovych et al., 2021]. CASP has driven advances by promoting rigorous data collection and innovation, leading to high-accuracy tools like AlphaFold2 and RosettaFold [Jumper et al., 2021, Baek et al., 2021]. Adopting a similar framework could significantly enhance the accuracy and reliability of genome annotation tools, improving our understanding of genome function.

Software benchmarks are subject to many caveats that are the result of inevitable limitations. In spite of these caveats, benchmarks serve a valuable purpose in providing an assessment of how tools perform on given datasets at a particular point in time. They can also identify performance limitations and directions where further improvements may be made. In brief, we: 1. use default parameters in each case to avoid inflating individual tool performance [Nießl et al., 2022], 2. we may apply some tools in a manner in which they were not designed (e.g. a transcriptome tool can be applied to genomic sequences), and 3. Some of the control data may be mislabelled, leading to conflicted outcomes. These caveats have been documented in detail elsewhere [Weber et al., 2019].

In this study, we evaluate the effectiveness of *de novo* protein-coding annotation tools of eukaryotic nucleotide sequences. These tools differentiate between nucleotide sequences or alignments that are either constrained by genetic codes or are non-coding. Given the distinct statistical signatures that coding sequences typically exhibit compared to non-coding ones, this task should be feasible using bioinformatic methods. The findings of this research have significant implications for the long non-coding RNA (lncRNA) research community, which is highly dependent on these tools to identify non-coding transcripts [Ponting and Haerty, 2022].

## Results

### Tool accuracy

Some clear trends have emerged from our analysis of coding sequence detection of reference datasets. The alignment-based tools RNAcode and PhyloCSF show a clear accuracy advantage over the single sequence approaches (Figure 1A-C, Supplementary Figures S1-S7). This ranking holds across each accuracy measure (Area Under ROC Curve (AUC), Sensitivity, Specificity and the Matthews Correlation Coefficient (MCC)) with RNAcode consistently outperforming PhyloCSF. The tcode tool, which is an implementation of a method proposed in 1982 is not only faster, but also significantly more accurate than modern sequence-based tools.

**Figure 1.**
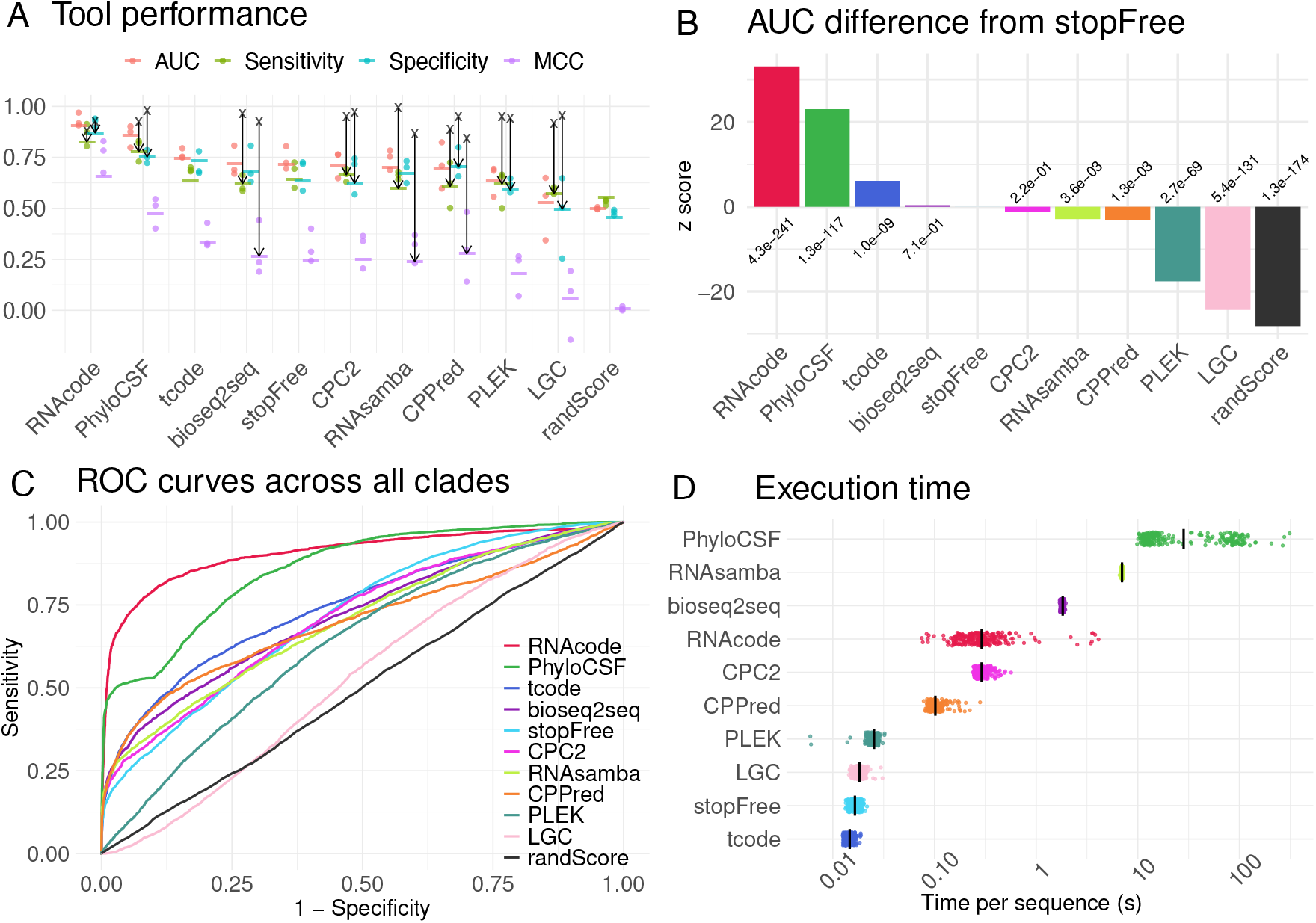
Coding potential tools, accuracy and speeds. **A:** A jitter plot showing the mean Area Under ROC Curve (AUC), Sensitivity, Specificity and Matthews Correlation Coefficient (MCC) values for each software tool and clade. Tools are ordered by the median AUC (mean performance values are indicated with a tick mark for each metric and tool). The black crosses and arrows indicate the accuracy metrics reported by the authors of each tool on their own test datasets. **B:** Bar plot showing Z-score differences of each tools ROC curves from our naive stopFree tool, with notated p-values. **C:** ROC curves showing the ability of each tool to discriminate between positive coding sequences and negative unannotated, length matched genome regions, and shuffled controls. The x-axis displays 1-Specificity (false-positive rate) and the y-axis displays the Sensitivity (true-positive rate). Each tool maps a trajectory from (0.0, 0.0) to (1.0, 1.0) through this plane as thresholds are lowered from the maximal to minimal value values for the tool on the test data. Ideal tools pass through the point (0.0, 1.0). **D:** A jitter plot showing the runtimes for each tool, measured in seconds. Runtimes were recorded by running each tool on sequences of varying lengths on an isolated computer.

We were particularly surprised to find that a naive tool, stopFree, that simply reports the length of the longest stop-codon-free region for a sequence, outperforms the widely used tools CPC2, PLEK, LGC and CPPred. This was a consistent finding across the performance metrics and datasets. To assess this further, we performed a permutation-based test to compare the ROC curves for each tool against the baseline stopFree tool (Figure 1B) [Venkatraman, 2000]. Three tools performed significantly better than the baseline stopFree tool: RNAcode (*z* score= 33.2, *p* = 4.3*x*10^−241^), PhyloCSF (*z* score= 23.1 *p* = 1.3*x*10^−117^), and 1982 sequence-based tool tcode (*z* score= 6.1, *p* = 1.01*x*10^−9^). The performances of bioseq2seq and CPC2 were not significantly better or worse than the stopFree tool. Whereas RNAsamba, CPPred, PLEK and LGC produced ROC curves that indicated a significantly worse performance than the stopFree baseline (*p <* 0.05 for each).

As a further assessment, we evaluated relative performance of tools on short open reading frames (sORFs). The results are relatively similar to the exonic results described above (Supplementary Figures S8&S9). The ranks of bioseq2seq and CPC2 climb to first and second/third (depending on accuracy metric), while PhyloCSF performance drops to below stopFree and LGC performs worse than the random number generator. Given the sORF lengths nest within the exon length distributions this shows the high dependence of bioseq2seq and CPC2 on the presence of in-frame start and stop codons (Supplementary Figures S9B).

### Self-assessment disparities

For most tools we note that there were large differences between the self-reported accuracies and the ones we calculated. Although this trend has previously been reported [Buchka et al., 2021], we were surprised by the degree of discrepancy (black arrows, Figure 1B & Supplementary Figure S1. The RNAcode publication reports the lowest self-reported accuracy statistics of all included tools and is yet the most consistent with our independent accuracy measurements (Figure 1A, Supplementary Figure S1,Supplementary Table S4). All self-reported accuracy statistics for the other tools have a much larger discrepancy with our results, which are ranked by degree of disparity in Supplementary Figure S1B.The possible causes of these discrepancies are discussed further in the conclusions.

### Computational efficiency of selected tools

In terms of computational efficiency, we found that the linear, composition and sequence-based tools tcode, stopFree, LGC and PLEK shared roughly equivalent runtimes of 0.01 − 0.03 seconds per sequence. CPPred, CPC2 and RNAcode had similar runtimes of 0.1 − 0.4 seconds per sequence, and bioseq2seq, RNAsamba and PhyloCSF are slower tools, needing 2.0 − 50 seconds per sequence (See Figure 1D, Supplementary Table S4). As can be seen in Supplementary Figure S2,most tools are relatively insensitive to sequence length at these length scales, with only the alignment-based methods showing a clear dependency between runtime and sequence length.

### Integrity of the control datasets

For most of the tools we tested, the coding potential scores for the two negative control sets (intergenic and shuffled sequences) of each clade (Animalia, Fungi, and Plantae) are generally very consistent (Supplementary Figures S4-S7), indicating that both shuffled and random intergenic regions can serve as valid negative controls for coding annotation tools, except two tools, RNAsamba and LGC.

RNAsamba scores show large differences between animal intergenic and shuffled sequences, with the shuffled scores being high and similar to the values for coding sequences. This suggests that RNAsamba identifies intergenic sequences rather than coding sequences when run on animal and plant derived sequences Supplementary Figure S6.

The LGC scores for plant genomes have a large discrepancies between the intergenic and shuffled negative controls, which may be due to the low C+G content in plant intergenic regions. These regions received higher scores than coding sequences, as shown in Supplementary Figure S5.This suggests a flaw in the LGC model’s ability to differentiate between non-coding and coding sequences with low C+G content [Wang et al., 2019].

### Relationship between citations and accuracy and speed

As observed in previous studies, citations rates do not reflect the accuracy or computational efficiency of bioinformatic tools [Gardner et al., 2022]. This lack of a relation can be confirmed here by comparing tool rankings from Figure 3B and Figure 1B In particular, RNAcode citations are approximately the same as two of the tools that performed barely better than a random number generator. In our hands RNAcode consistently outperforms competing tools. Meanwhile, the most highly cited, and presumably most widely used tool CPC2, performed no better than the baseline stopFree scores while being one of the least computationally efficient sequence based tools (Figures 3B & 1D).

## Discussion

### Evolutionary vs single sequence tools

Our evaluation shows that tools using evolutionary conservation patterns, like RNAcode and PhyloCSF, are more accurate than those based on single sequences alone. As we continue to expand the genomic tree of life [Christmas et al., 2023, Kuderna et al., 2023], the usefulness of deploying such tools across different clades will increase significantly. These tools benefit from incorporating variations in nucleotide sequences that preserve protein functions, such as synonymous changes, conservative amino acid substitutions and frame-preserving insertions and deletions. This approach not only enhances accuracy but also supports the ‘selected effect’ definition of functional elements [Linquist et al., 2020].

The only sequence based tool, tcode, to outperform the naive stopFree score is faster and more accurate that modern sequence-based tools. These three tools (RNAcode, PhyloCSF and tcode), published in 2011, 2011, and 1982 respectively, highlight how little genuine progress in the field has been made in recent years.

### Challenges and recommendations for future development

This study adhered to key principles that should guide future tool development and evaluation. This is in contrast with some current tools that overlook essential aspects of mathematical modeling such as understanding the data and its applications. We highlight three main areas for consideration:

1. *Sampling controls:* We used well-annotated reference eukaryotic genomes outside any training dataset for our positive controls, and question the need for species-specific tools. Given the conservation of the genetic code and translation machinery [Petrov et al., 2015, Shulgina and Eddy, 2021], more generalized coding detection tools are feasible. Moreover, coding detection tools must be robust to biological and technical deviations from curated data, including truncated sequences, splicing errors, frame-shifting elements, recoding, non-AUG starts and non-standard genetic codes [Steijger et al., 2013, Pickrell et al., 2010, Dvinge and Bradley, 2015, Andreev et al., 2022, Rodnina et al., 2020, Shulgina and Eddy, 2021].
2. *Dataset construction:* Both positive and negative datasets must be challenging and well-matched to control confounding factors such as sequence length and C+G content. This ensures that coding sequence scores will be robust and distinct from genomic background. Using short and long ncRNAs as negative controls is insufficient (see below).
3. *Dataset representation:* Evaluation datasets should mirror the typical genomic composition, which for many species includes a predominance of junk and functional non-coding sequences. Evaluation metrics must be robust to the presence of imbalanced datasets [Whalen et al., 2022].

In this study, several widely used tools performed poorly in distinguishing coding from non-coding sequences, illustrating once more that software popularity does not equate to accuracy or efficiency [Gardner et al., 2022]. Despite high self-reported accuracy, our independent evaluations revealed large discrepancy with measurements, suggesting the need for more rigorous validation measures (See Figure 1A and Supplementary Figure S1)[Buchka et al., 2021].

### Cease relying on highly curated datasets for training and testing

The meticulously curated mRNA sequences from RefSeq and GENCODE [Frankish et al., 2021, Rajput et al., 2019] are, in our view, unsuitable as positive controls for training or testing coding annotation tools. These datasets have undergone extensive refinement over decades by many curators who incorporate additional experimental data such as Ribo-seq and mass-spec to refine annotations [O’Leary et al., 2016, Sayers et al., 2024]. However, typical real-world datasets such as new genome and transcriptome assemblies contain errors such as truncations, sequencing and assembly problems that cause frame-shifts and mis-spliced transcripts. Consequently, highly curated datasets are a trivial problem set that exaggerates the significance of inherent sequence features, such as the longest canonical ORFs. This issue is illustrated by the discrepant results between the exon- and sORF-derived datasets, with some tool performances improving markedly, despite the only major difference being the presence of in-frame start and stop codons. This is likely the cause of the many misclassifications by many tools in our practical assessment scenarios.

The problematic selection of negative training datasets is a further issue. We noted that either short or long ncRNAs have been used as negative datasets for training some of the above tools (see tools descriptions in the Supplementary Materials). Rfam-derived datasets are an inadequate negative control for this problem as these short non-coding RNAs have profoundly different lengths and sequence compositions compared to coding and typical genomic sequences. Furthermore, the majority of lncRNAs have been labelled lncRNAs because they have (a) some evidence of transcription, and (b) lack an obvious open reading frame [Xu et al., 2017]. Therefore, lncRNAs have shorter ORFs than random sequences and therefore are a trivial and inadequate negative control for training or testing a coding-potential prediction tool. Finally, lncRNAs have a very different length distribution relative to mRNAs, therefore trivial tools could distinguish between these coding and non-coding using trivial length-associated measures.

In short, the tools that underperform here excel at solving a non-existent problem in genomics: distinguishing error-free, curated coding sequences from shorter non-coding RNA sequences in model organisms with up to 99.85% accuracy.

### Class balance

We noted that many tools trained and evaluated their performance on balanced classes of negative and positive datasets. In reality the ratio of coding to non-coding is likely to be low in eukaryotes (approximately 1 to 100 the human genome for example). In order to evaluate performance under realistic conditions we recommend that realistic positive to negative class ratios are maintained. Assessments should use metrics that are robust to class imbalance, rather than artificially attempt to balance positive and negative classes [Whalen et al., 2022, McDermott et al., 2024].

### Limit sequence similarity between test and training

a standard practice for applying machine learning techniques in computational biology is to ensure test and validation sets have limited homology or similarity to the training set [Söding and Remmert, 2011, Walsh et al., 2016, Petti and Eddy, 2022]. Many of the tools assessed here did not report controlling for similarity between their training and test sets (See “tool descriptions” in the Supplementary Information), as a consequence are likely to have artificially inflated the performance measures for their tool ().

### Study limitations

Firstly, this study is very eukaryote-specific, we have not assessed tools on more diverse genomic data from organelles, protists, archaeal, bacterial or viral sequences. Species that utilise novel genetic codes may prove to be a particularly challenging cohort for assessments [Shulgina and Eddy, 2021].

Secondly, we have used a relatively balanced positive and negative datasets (ratios of approximately 1:3). On more representative datasets the negative datasets vastly outnumber the positive (e.g. approximately 99% of the genome is non-coding in human) [Frankish et al., 2023]. So, ideally approximately 30-50 times more negative as positive sequences should be used to reflect realistic coding:non-coding ratios [Whalen et al., 2022].

Thirdly, we have “abused” some tools by not applying them to the scenarios for which they were developed. For example, transcriptome classification tools expect positive stranded and full-length ORFs in sequences that are free of introns. Also, our intergenic negative control regions are likely to be less well conserved than coding sequence. However, the alignment depths and percent-identity are similar (Table S3), whereas the shuffled alignments have identical depth and identity. We found there was little difference in score distributions between intergenic and shuffled for the alignment-based tool results, as a consequence the alignments for intergenic sequence appear to be valid negative controls.

Finally, we have not generated more complex sequence sets for evaluation. We did not attempt to assemble full length transcripts or simulate sequencing and assembly errors found in typical genome and transcriptome datasets. This may be a fruitful avenue for future studies.

### Future efforts

We have highlighted the scope for further development of tools to address the coding sequence annotation challenge. We note that the existing tools have not employed some of the latest machine-learning methods, nor do they include more complex information for conserved peptide predictions, for example the increased covariation in coding sequence alignments has been observed in recent analyses [Cooper and Gardner, 2020, Gao et al., 2023]. We also note there has not been much work in identifying the strengths of the statistical features that contribute to predictions.

From a biocuration perspective, there has been relatively little development of the “conservome” for coding sequences, i.e. large-scale screens of conserved sequence elements with characteristic signatures of evolutionarily conserved coding sequences [Mudge et al., 2019]. These screens may prove vital for identifying functional alternative splicing, alternate start or stop codons, short open reading frames and other conserved non-canonical protein-coding sequences.

Finally, the results presented here further highlight the need for well maintained, robust, up-to-date and accurate software. To be achieved this will require further support for long-term software maintenance and career incentives that encourage these activities [Siepel, 2019].

## Materials and Methods

Our methodology is divided into these key sections: (1) Dataset preparation, where we describe the acquisition and preparation of coding and non-coding sequences used as controls; (2) Performance measures, where we explain the methods used to analyze the results and verify the significance of our findings. (3) Tool inclusion, detailing the criteria for selecting the annotation tools evaluated in this study; and (4) Benchmarking Strategy, outlining the protocols for tool performance assessment, including accuracy, efficiency, and computational demand; These ensure a comprehensive evaluation of each tools’ capabilities in distinguishing between coding from non-coding genomic sequences.

### Data selection: positive & negative control sequences

To improve the assessment of software predictions for protein-coding nucleotide sequences and mitigate biases, we avoid commonly used reference genomes such as *Homo sapiens* and *Mus musculus*. We selected representative species from three eukaryotic clades (mammals, plants, and fungi) that have not previously been used in software training. These are *Felis catus* (house cat), *Cucumis melo* (melon), and *Aspergillus puulaauensis*. Further details are available in Supplementary Tables S2 and S3 and Figure 2.

**Figure 2.**
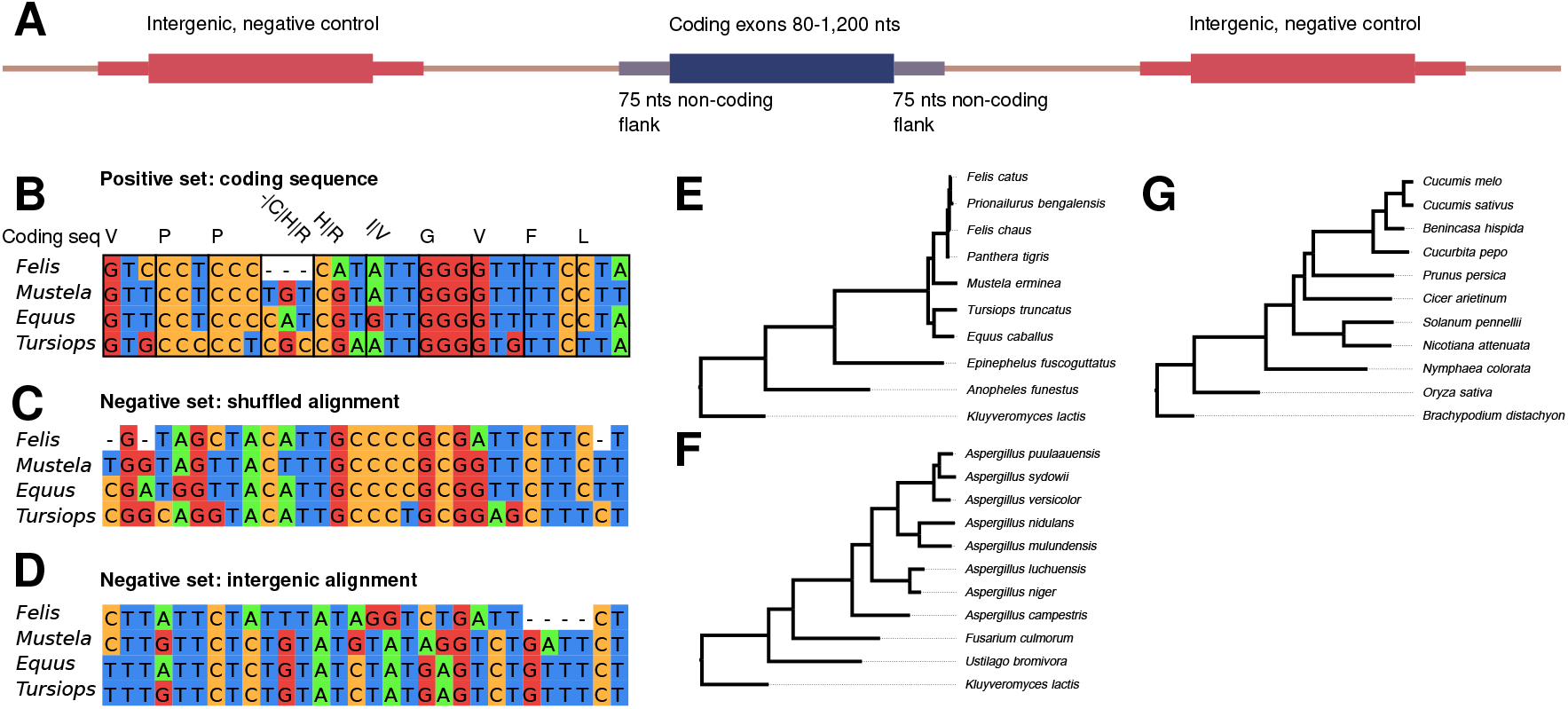
Controls for benchmarking protein-coding detection tools. **A:** Annotated exons of 80 to 1,200 nucleotides (nts) are selected from reference genomes, flanking 75 nts are also included with these positive control sequences. These are length-matched with genomically nearby intergenic sequences which serve as one source of negative controls. Shuffled coding sequences are used as a further source of negative control. **B-D:** Exemplar multiple sequence alignments, these include a partial exon (**B**), with corresponding negative controls from a shuffled alignment (**C**) and a neighbouring intergenic region (**D**). For alignment **B** the corresponding amino-acid(s) of each codon is shown along the top line. **E-G:** Phylogenetic trees for genomic reference and selected related genomes for producing alignments, these are the (**E**) animalia (reference: *Felis catus*), (**F**) fungi (reference: *Aspergillus puulaauensis*), and (**G**) plants (reference: *Cucumis melo*).

Up to ten further genomes that span a range of phylogenetic distances are selected for each group, these are used to construct multiple sequence alignments for the comparative tools (Table 1). Each selected genome assembly has a high BUSCO completeness score [Simaõ et al., 2015, Manni et al., 2021] (Supplementary Table S3), and have numbers of annotated coding genes that is consistent with related genomes.

**Table 1.**
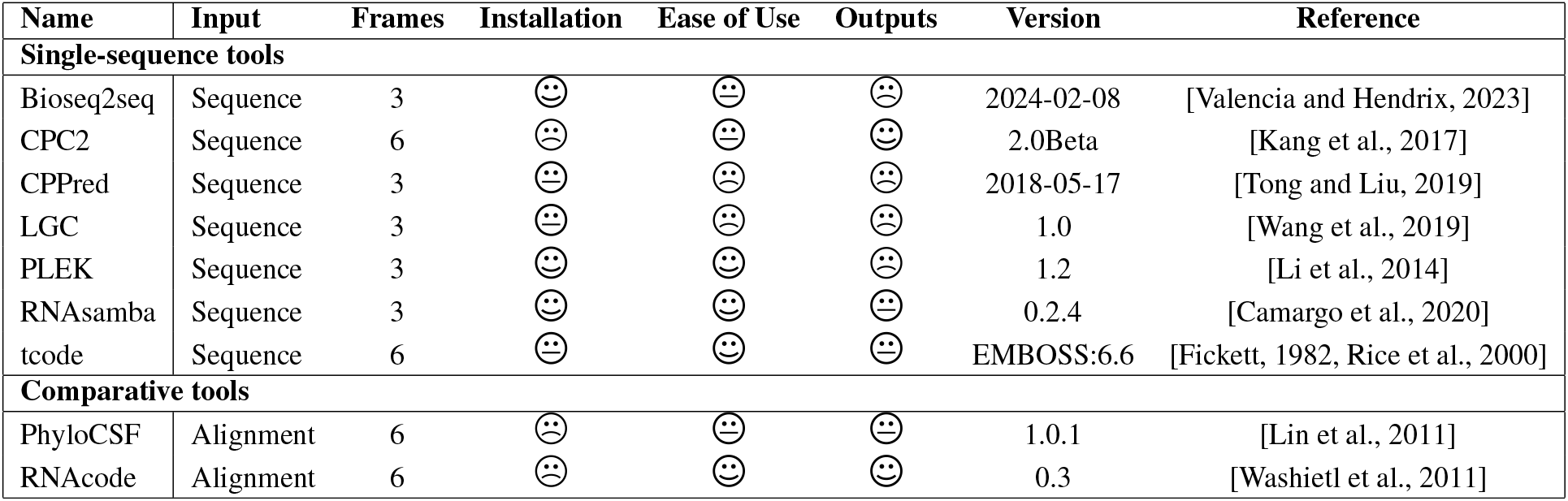
Summary of assessed tools. We have annotated the type of input each tool requires, the number of frames they scan (frames 1-3 [fwd], or frames 1-6 [fwd+rev]), the version and references. In addition, tools have been assessed for their ease of installation on a modern *nix environment, and for their ease of use, and whether the outputs included useful information like ORF coordinates as well as scores. The categories for each were: Excellent: 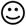, Acceptable: 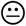, Unsatisfactory: 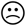. Further details on these assessments can be found in **Supplementary Text**.

The positive control coding sequences are derived from GenBank annotations. We collapsed overlapping annotations and randomly sampling 1,200 coding exons and 40 sORFs from each reference genome. Further realism is introduced into the test dataset by including 75 nucleotides of intron or UTR from both sides of each exons (refer to Figure 2A). Negative control intergenic sequences of the same lengths are selected from adjacent upstream and downstream intergenic regions that exceed one kilobase (kb) in length (for summary statistics see Supplementary Table S3). The intergenic sequences are compared with protein in the UniProt database and possible coding sequences (e-value *<* 10^−10^, bitscore *>* 31) were removed using translated searches with MMseqs2 (v15.6f452) [Steinegger and Söding, 2017].

Alignments for comparative tools were made for intergenic and coding sequences (Figure 2). Reference sequences are queried with MMseqs2, and top-scoring matches for each genome selected for multiple sequence alignment with Clustal Omega (1.2.4) [Sievers et al., 2011]. A further negative control set is produced by shuffling coding alignments with `esl-shuffle` (Easel 0.48 (Nov 2020)).

Each of the selected prediction tools were run on both the coding and non-coding sequences or alignments. Subsequently, the optimal coding potential scores for each input for each tool were used as a basis for computing performance statistics, as defined in the below ‘Performance metrics’ section. For the tools that only scan the forward strand, the reverse complement of each input sequence was also screened. In instances where multiple scores are returned for a sequence (in any of the six-frames), the best score is selected (highest or lowest, depending on which is a better indicator for a coding sequence).

### Performance metrics and output management

Sequences are ranked by prediction score, and a sliding threshold is used. The coding sequences that score at or above a threshold are labelled “true positives” (TP), negative control sequences at or above the threshold are “false positives” (FP), coding sequences below a threshold are “false negatives” (FN), and negative control sequences below a threshold are “true negatives” (TN). An optimal score threshold is selected for each tool based on the minimal distance to the [0.0,1.0] coordinate on the ROC plot (i.e. “closest.topleft” in the `R pROC` package). This best balances the sensitivity and specificity for each tool [Robin et al., 2011].

For each possible score threshold the *sensitivity, specificity* and *Matthews correlation coefficient (MCC)* are computed, these are defined below. A “receiver-operator characteristic” (ROC) plot is generated for each tool (Figure 1A) using the pROC package (version 1.18.5) [Robin et al., 2011] and the “AUC” or Area Under the ROC curve computed for each tool, which we note has a theoretical range [0.5, 1.0], and is robust to potential issues of class imbalance [McDermott et al., 2024].

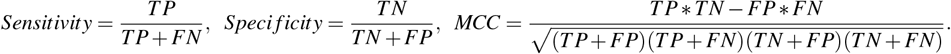

The runtimes for each tool were collected on a standard, isolated desktop computer (Intel 12-core processor i9-8950HK, 32GB RAM) running a Linux operating system (Ubuntu 22.04.2 LTS, Linux kernel 6.2.0-36). We recorded times for predictions on 240 sequences with lengths ranging from 232 to 1114 nts. The three-frame, top-strand only tools were run on both the forward, and reverse complement sequences. The average runtimes for each tool can be viewed in Figure 1D, the dependency between sequence length and runtime can be seen in Supplementary Figure S2, average values can be viewed in Supplementary Table S4, column O.

#### Tool inclusion criteria

In order to select tools for inclusion in the benchmark we iteratively searched the published literature (GoogleScholar, PubMed), examining previous benchmarks, software tool papers and reviews of the topic [Fickett and Tung, 1992, Li et al., 2020, Zheng et al., 2021, Singh and Roy, 2022]. We extracted candidate tools from the text and from the reference sections, these have been assessed against the following inclusion criteria: 1. The primary purpose of the tool is to predict the protein-coding potential of eukaryotic nucleotide sequences or alignments, 2. The tool is publicly accessible, 3. The tool is able to be installed and executed, 4. The tool is generalised, and not restricted to a single species or phylogenetic group, 5. The tool is unique, i.e. is not a clone or slight modification of an existing tool, 6. The tool is **not** based on homology to known proteins, 7. The tool has been used recently, and there is evidence that it is or may be popular.

The tools that we assessed against these criteria are listed in Supplementary Table S1. A total of 36 tools were assessed and **75% (27)** did not meet at least one of the above criteria. The most common reasons for excluding a tool was the installation was unnecessarily complex or failed on a modern linux computing environment (criterion #3, 41% failed, 11/27), the tool was not generalised (criterion #4, 33%, 9/27) or was not publicly accessible (criterion #2, 26%, 9/27) (Figure 3A, Supplementary Table S1).

**Figure 3.**
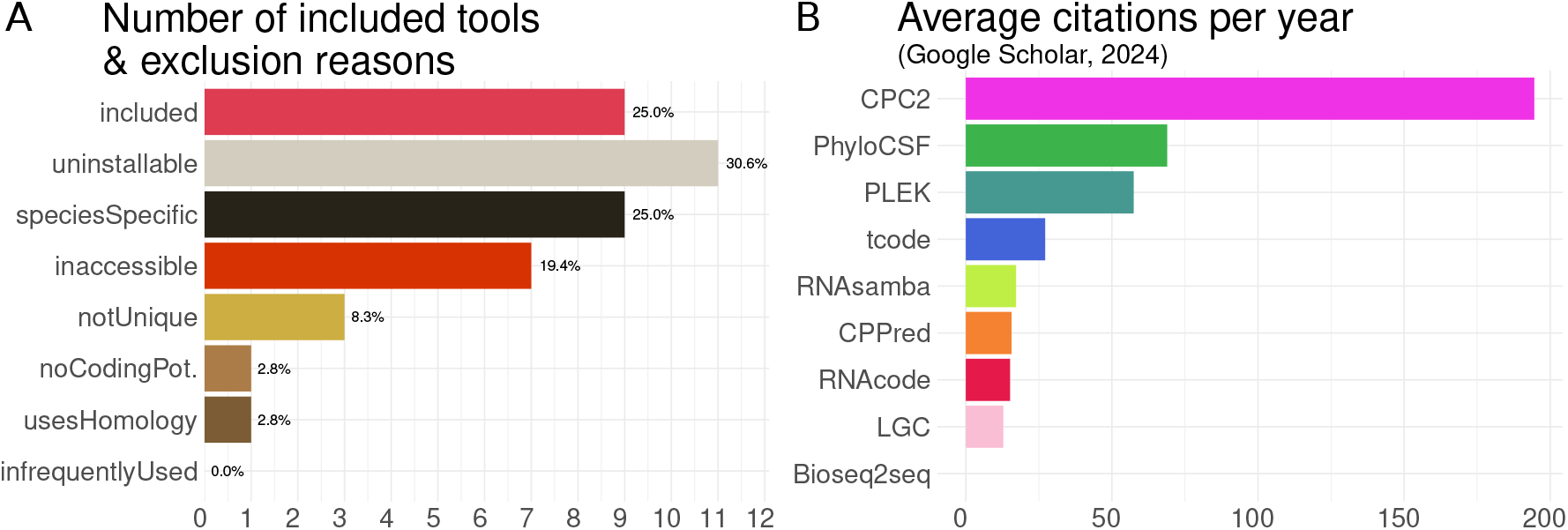
Software inclusion, exclusion and popularity. **A:** The proportions and counts of candidate software tools (total of 36), and the reasons some tools were excluded. See text and Supplementary Table S1 for further details. **B:** The average number of citations per year for each of the included tools (citation counts were collected in March 2024 from Google Scholar).

### Brief tool descriptions

In the following section we briefly describe each of the software tools that met the above inclusion criteria for this study. See Supplementary Information for detailed information on sources of training and test data, the number of sequences used as positive and negative controls for each, whether summary statistics for each (e.g. mean length and C+G content) are reported and whether similarity between training and test datasets were accounted for.

#### Base-line tools

The authors of this manuscript added a naive coding potential finder “`stopFree`”, and employed a random number generator “`randScore`”. Both tools serve as base-line scores for a minimal and lowest expected performance in our evaluation that are free of any machine-learning based training. The simplest coding potential measure we have identified is the length of the longest in-frame stop-free region for any input sequence (i.e. the longest run of `UAA`, `UAG`, and `UGA` free sequences over each of the six possible reading frames). This value conforms with the most general definition of an ORF [Sieber et al., 2018].

We expect the stopFree measure to be robust to common sequence errors and non-canonical features such as truncations, non-AUG start codons [Andreev et al., 2022], mis-splicing, and many forms of sequence error [Steijger et al., 2013, Rodnina et al., 2020]. We anticipate that stopFree as a minimal, single-metric tool will serve as a lower bound on tool accuracy for more sophisticated tools.

Meanwhile, randScore uses a random number generator that generates integers in the range [1, 100] for any input sequence, and is expected to be the worst possible prediction tool (also known as “monkey with a pencil”).

#### Protein-coding potential tools

Various approaches were employed by the tools that were benchmarked. We briefly describe them here, while more detailed tool descriptions, discussion of their training and test datasets, sequence/alignment features, command-line parameters, installation, user experience and output notes can be found in the Supplementary Information.

For the sequence-based tools, `Bioseq2seq` [Valencia and Hendrix, 2023] and `RNAsamba` [Camargo et al., 2020] use neural network models. `CPC2` “Coding Potential Calculator 2” [Kang et al., 2017], `CPPred` “Coding Potential PREDiction” [Tong and Liu, 2019] and `PLEK` [Li et al., 2014] utilise support vector machines. `LGC` “ORF Length and GC content” [Wang et al., 2019] employs a maximum likelihood method. `tcode` [Fickett, 1982, Rice et al., 2000] uses an implementation of the Fickett TESTCODE statistic which is a probabilistic method.

For the alignment-based tools, `PhyloCSF` “Phylogenetic Codon Substitution Frequencies” [Lin et al., 2011] is a maximum likelihood method, while `RNAcode` [Washietl et al., 2011] is a comparative analysis tool.

## Author engagement

In order to ensure methods are used correctly, a preprint of these results and methods was shared with corresponding authors for each tool. We have elected to do this in order to minimise disagreements between benchmark and method authors, and ensure that we have adequately described, and not misrepresented any tools.

## Declarations

### Ethics approval and consent to participate

Not applicable

### Consent for publication

Not applicable

### Availability of data and materials

A slim github repository for researchers who wish to use just the tool prediction scores, control sequences and alignment is available here:

https://github.com/Gardner-BinfLab/PCPBSlim/

The full github repository required to replicate the genome sequence sampling, shuffling and multiple sequence alignment, the scripts to run each tool, and calculate summary accuracy statistics is here:

https://github.com/Gardner-BinfLab/PCPBFull/

## Authors’ contributions

this work was conceived by DJC, THC, ST & PPG, designed by DJC & PPG, the analysis was carried out by DJC, PPG & MAB, interpretation of the results was by DJC & PPG, the manuscript was drafted by DJC & PPG. All authors revised and edited the final manuscript. All authors approve the submitted version.

## Acknowledgements

We have received valuable feedback from the Genomics Aotearoa research community. Professor Mercelo Carazzolle and Dr. Antonio Camargo for feedback on their tool, `RNAsamba`.

