## Supplementary text and figures S1-S9 for "Evaluating computational tools for protein-coding sequence detection: Are they up to the task?"

**Abstract**

**Keywords**
Genome annotation, transcriptome, protein coding, open reading frame, ORF, benchmark

[1] *Department of Biochemistry, University of Otago, Dunedin, New Zealand.*
[2] *The New Zealand Institute for Plant and Food Research Limited, Lincoln, New Zealand.*

### Supplementary information

**References** **15**

#### S1.1 Detailed tool descriptions, command line parameters and installation, running and user-friendly status

In the following section we describe each of the software tools that met the inclusion criteria for this study. We have systematically identified where possible:

- the sources of training and test data
- the number of sequences used as positive and negative controls for each
- whether summary statistics for each (e.g. mean length and C+G content) are reported
- whether similarity between training and test datasets were accounted for
- self-reported accuracy statistics for each tool (sources are provided in Supplementary Table S4)
- whether or not all six frames are assessed by the tool and
- whether or not coordinates of predicted ORFs are reported.
- the command-line parameters
- justify the Excellent (🙂), Acceptable (😐) or Unsatisfactory (🙁) classifications for the installation, user experience and output of each tool (Table **??**)
- Installation:
  - Is the tool well documented?
  - Is the tool current?
  - Are dependencies/other tool installations described?
  - Is package management used?
  - Where were the installation instructions? E.g. a text file, github, or other.
- User experience:
  - Does the tool have well-documented use cases?
  - Are tool options clearly documented?
  - Did the tool crash frequently?
  - Are there verbosity options that provide useful dialogue?
  - Were informative warnings printed?
  - Were examples provided?
- Output:
  - Was the output formatted in an easy-to-parse flat file (e.g. tsv, csv, gff, bed, . . . )
  - Was useful information provided (e.g. coding score/probability, ORF coordinates, ORF strand)
  - Was a complex wrapper required to screen the output?

##### S1.1.1 Base-line tools

**stopFree** and **randScore**: The authors of this manuscript constructed a naive coding potential finder, and employed a random number generator. Both tools serve as base-line scores for a minimal and lowest expected performance in our evaluation. The simplest coding potential measure we identified is the length of the longest in-frame stop-free region for any input sequence (i.e. the maximum run of `UAA`, `UAG`, and `UGA` free sequences over each of the six possible reading frames). This conforms with the most general definition of an ORF as discussed by Sieber *et al.* [1].

We expect the stopFree measure to be robust to common sequence errors and non-canonical features such as truncations, non-AUG start codons [2], mis-splicing, and many forms of sequence error [3, 4]. We also note that no machine learning methods and no training data were used in the development of this tool, and as far as we are aware this is the first assessment of this metric. We anticipate that stopFree as a minimal, single-metric tool that will serve as a lower bound on tool accuracy for more sophisticated tools.

Meanwhile, randScore uses a random number generator that uses the integers in the range $[1,100]$ for any input sequence, and is expected to be the worst possible prediction tool (also known as "monkey with a pencil").

The command-line parameters used for this evaluation were:

```
$ python3 stopFree input.fa outputFile

$ randScore=$((1 + $RANDOM % 100)); echo $randScore  > outputFile
```

###### S1.1.2 Sequence-based tools:

###### S1.1.3 Bioseq2seq

**Bioseq2seq** is a neural network model of nucleotide to protein translation [5]. It is unclear from the bioseq2seq paper what and how many specific variables are extracted from each sequence, but we understand that a Fourier transform is used to capture potential sequence periodicity signals. The training data is derived from RefSeq (release 200) transcript and protein sequences derived from eight mammalian species (*Bos taurus*, *Pan troglodytes*, *Gorilla gorilla*, *H. sapiens*, *Macaca mulatta*, *M. musculus*, *Pongo abelii*, *Rattus rattus*). From 63,272 lncRNA and mRNA transcripts $< 1,200$ nts training, validation and test sets were sampled, using an 80/10/10 proportion for each. The similarity between the sets was restricted using CD-HIT-EST-2D, BLAST and Needleman-Wunsch alignment to exclude transcripts $> 80\%$ nucleotide sequence similarity. The test set contained 1,913 lncRNAs and 1,791 mRNAs, approximately a 1:1 ratio. The length and C+G contents for both the positive and negative datasets is not mentioned. The self-reported overall accuracy statistics for bioseq2seq are an F1 of 0.961, a sensitivity (recall) of 0.963, a PPV (precision) of 0.960, an MCC of 0.925 and AUPRC of 0.994 [5]. We further note that only the forward strand (frames 1, 2 and 3) are evaluated by this tool, and the coordinates of predicted ORFs are not provided in the output.

The command-line parameters used for this evaluation were:

```
$ python translate.py --checkpoint best_bioseq2seq-wt_LFN_mammalian_200-1200.pt
                      --input input.fa         --output outputFile
$ python translate.py --checkpoint best_bioseq2seq-wt_LFN_mammalian_200-1200.pt
                      --input input-REVCOMP.fa --output outputFile
```

Documentation for the install process located on github is simple and effective. Installation requires creating a Python virtual environment, using 'pip' with a requirements file and executing a bash install script. This ensures all dependencies are correctly versioned and not in conflict with installed packages. Installation is excellent. ☺

Usage documentation is clear and includes examples, making it easy to get started. However, it falls short in explaining the function of all parameters. Additionally, there are no simple warnings for user errors; it only throws general Python errors for missing or incorrect parameters. User experience is acceptable. 😐

Plain text output files are not formatted in an easy to parse manner. Metrics are distributed over multiple lines and have inconsistent spacing and colons as delimiters. There are only two useful metrics provided, coding probability and coding verdict. Output is unsatisfactory. ☹

###### S1.1.4 CPC2

**CPC2** "Coding Potential Calculator 2" is an extension of CPC1 [6]. A random forest was used to select informative intrinsic features coding features (e.g. Fickett score [7], canonical ORF length, "ORF integrity" and isoelectric point). These features were used to train a support vector machine on approximately 18,000 high confidence *Homo sapiens* coding transcripts and 10,000 non-coding transcripts. The source of these transcripts is unclear from the manuscript since the numbers do not match the cited data source [6]. An "independent" test set of mRNAs was generated by intersecting the well curated Swiss-Prot and Refseq databases from *Homo sapiens*, *Mus musculus*, *Danio rerio*, *Drosphila melanogaster*, *Caenorhabditis elegans* and *Arabidopsis thaliana*. Non-coding transcripts were sourced from Ensembl and EnsemblPlants. Ratios between the positive and negative datasets ranged from approximately 1:2 to 4:1 across the species. Sequence similarity between the coding sequences was reduced to 90% sequence identity threshold with CD-hit. It is unclear from the methods section whether sequences homologous to the training dataset where removed from the independent test set. The self-reported overall accuracy statistic for CPC2 was 0.96, with specificity 0.97 and sensitivity of 0.95 [6]. Characteristic summary statistics for the positive and negative datasets such as length and C+G content were not reported. We further note that both the forward strand (frames 1, 2 and 3) and reverse (frames 4, 5 and 6) are evaluated by this tool.

The command-line parameters used for this evaluation were:

```
$ python2.7 CPC2.py  -r --ORF -i input.fa -o outputFile
```

Installation instructions available on their website are slightly confusing. The website download page details two options for installing. Method one is called "standalone", which contains the filename "beta" and is based on python2.7. Method two which supports Python3 but has the filename "standalone". The rest of the instructions pertain to the "standalone" version, which is seemingly for the python2.7 version "beta" and not the python3 version "standalone". Biopython is required, but no version is stated. Installation is unsatisfactory. ☹

Documentation for running the tool consists of a single example run. The only options documented are found by running the tool without parameters. Incorrect usage results in useful error messages. User experience is acceptable. 😐

Output is provided in simple plain text and tab delimited format with headers and is easy to read and parse. Output includes important metrics such as coding probability, ORF start and Fickett score, as well as transcript and peptide lengths. Output is excellent. ☺

##### S1.1.5 CPPred

**CPPred** "Coding Potential PREDiction" is a Support Vector Machine (SVM) based coding prediction tool [8]. Thirty eight sequence features were ranked and selected using "mRMR-IFS" [9], these included the features used by CPC2 above, as well as hexamer scores, Gravy and protein instability metrics [10], and 30 "CTD" (composition-transition-distribution) features which appear to be mono- and di- nucleotide features, together with 20 "D descriptors" the definitions of which were unclear from the manuscript. A *H. sapiens* specific and an "integrated" model were trained on either *H. sapiens* alone, or *D. rerio*, *D. melanogaster*, *S. cerevisiae*, *C. elegans* and *A. thaliana* data. Positive coding sequences were obtained from NCBI RefSeq, while negative sets of sequences were based on ncRNAs from ENSEMBL release 90. We sampled the *H. sapiens* training datasets and found that the coding sequences had an average length of $\sim 3,800$ nts, while the average ncRNA length was $\sim 800$ nts. The C+G content of both sets is $\sim 47\%$. In total $\sim 50,000$ coding and $\sim 36,000$ ncRNA sequences were collected ($\sim 5:4$ ratio). These were randomly assigned to either a training set (two thirds) or a test set (one third). Sequence similarity between the test and training sets was reduced to a 80% sequence identity threshold with CD-hit. The self-reported performance metrics of the integrated model on integrated test data was 0.92 MCC, 0.99 AUC, 0.96 accuracy, 0.97 sensitivity and 0.95 specificity [8]. We further note that only the forward strand (frames 1, 2 and 3) are evaluated by this tool, and the coordinates of predicted ORFs are not provided in the output.

The command-line parameters used for this evaluation were:

```
$ python2.7 CPPred.py  -i input.fa         -hex Integrated_Hexamer.tsv
                       -r Integrated.range -mol Integrated.model
                       -spe Integrated -o outputFile
$ python2.7 CPPred.py  -i input-REVCOMP.fa -hex Integrated_Hexamer.tsv
                       -r Integrated.range -mol Integrated.model
                       -spe Integrated -o outputFile
```

Installation instructions are available on their website. While the version of biopython is provided, the python2.7 dependency is not listed. Installation is acceptable. 😐

Usage documentation consists of providing a template example as well as multiple other examples with differing option variables. The program command can become overly complex as it requires six command line options to run. It is also not possible to run multiple instances at once due to hard coded temporary file names rather than randomised temporary filenames. The absence of error messages made troubleshooting difficult. User experience is unsatisfactory. ☹

The output files include over 40 data fields yet lack important information such as predicted start or stop sites and in which frame the provided score is found. The output file is tab-delimited, and easy to parse, however there is no documentation defining each output metric. Output is unsatisfactory. ☹

##### S1.1.6 LGC

**LGC** "ORF Length and GC content" employs a maximum likelihood method to distinguish coding sequences from lncRNAs [11]. A polynomial is fit to the relationship between GC content and length of the top three ORFs ($> 100$ nts), parameters are estimated that minimise the root mean square error based on mRNA and lncRNA sequences from *H. sapiens*, *M. musculus*, *C. elegans*, *Oryza sativa* and *Solanum lycopersicum*. *H. sapiens* and *M. musculus* were obtained from RefSeq, lncRNAs from GENCODE. For the remaining species both sequence types were obtained from ENSEMBL, excluding ncRNAs $< 200$ nts. A total of $\sim 226,000$ protein-coding and $\sim 52,000$ lncRNA sequences were used in training (i.e. a ratio of coding positive to negative sequences of roughly 4 to 1). The summary statistics such as length and C+G content were not directly reported. The robustness of the approach is evaluated using all the plant, invertebrate, vertebrate and mammalian protein coding and ncRNAs from RefSeq (no version number was given, it is unclear if the mammal set is included within vertebrates too). Sequence similarity between the training and test sets was not discussed. The self-reported average performance metrics over six species for LGC were 0.94 accuracy, 0.92 sensitivity and 0.95 specificity [11]. We further note that only the forward strand (frames 1, 2 and 3) are evaluated by this tool, and the coordinates of predicted ORFs are not provided in the output.

The command-line parameters used for this evaluation were:

```
$ python2.7 LGC-1.0.py input.fa         outputFile
$ python2.7 LGC-1.0.py input-REVCOMP.fa outputFile
```

Installation documentation provided on their website is simple, but incomplete and out of date. Python 2.7 and biopython are required, however biopython now uses Python 3. The link to instructions on installing biopython is also no longer valid. The user needs to determine which version to install. Installation is acceptable. 😐

Usage documentation consists of providing one example. Command line options are non-existent with only an input and output being available. It is also not possible to run more than one instance of the software at a time due to non-random temporary file naming conventions. User experience is unsatisfactory. ☹

For output, documentation has simple descriptions of each of the output metrics. The output usefulness is hampered by not having any frame data or start and stop locations. The file is tab delimited and easy to collate data from, but unnecessarily repeats the documentation's explanations of each of the output fields which increased the visual clutter of the output. While the majority of scores ranged between -10 and 10, sometimes an output of -10,000 is given. Output is unsatisfactory. ☹

###### S1.1.7 PLEK

**PLEK** "Predictor of long non-coding RNAs and messenger RNAs based on an improved k-mer scheme" uses a SVM to discriminate lncRNAs from protein-coding mRNAs [12]. PLEK uses 1,364 k-mer weighted frequency features for $k$ values ranging from one to five (i.e. $\sum_{k=1}^{5} 4^k$ features). A SVM was trained on these features, with randomly selected 22,389 protein-coding transcripts from a *H. sapiens* RefSeq dataset (release 60) as a positive-set, and 22,389 *H. sapiens* long non-coding transcripts from GENCODE v17 formed a negative-set. In order to make the tool more robust to INDEL errors, INDELS of length zero to three were simulated with a rate ranging from 0% to 3%. A test set was generated by collecting novel transcriptome data for *H. sapiens* cell-lines MCF-7 and HeLa S3, the resulting sequences were allocated to either coding or non-coding sets by sequence alignment with RefSeq and GENCODE. There were 3,185 mRNAs and 121 lncRNAs corresponding to the MCF-7 data, and 3,045 mRNAs and 53 lncRNAs corresponding to the HeLa data. The potential for similarity between the training and test datasets was not discussed, and summary statistics such as length and C+G content were not reported, other than a limit $>$ 200 nt being used for each dataset. The self-reported average performance metrics over the two cell-line datasets were 0.947 and 0.955 sensitivity, and 0.958 and 0.925 specificity [12]. We further note that only the forward strand (frames 1, 2 and 3) are evaluated by this tool, and the coordinates of predicted ORFs are not provided in the output.

The command-line parameters used for this evaluation were:

```
$ python2.7 PLEK.py -fasta input.fa         -out  outputFile -thread #cpus
$ python2.7 PLEK.py -fasta input-REVCOMP.fa -out  outputFile -thread #cpus
```

Installation documentation on their website provides sufficient information to install and details prerequisites. Installation is excellent. ☺

Usage documentation is easy to follow and provides multiple examples as well as option parameter explanations. It works well with having multiple instances running and each can also be run with multiple cores using command line options. Erroneous usage produces useful error messages. User experience is excellent. ☺

Documentation provides no information on output files, however the output is a simple one-line text file. The only details in the output file are a coding verdict, followed by a score and the name of the sequence. The output usefulness is hampered by not having any frame or start and stop locations. The file is tab-delimited and easy to collate data from. Output is unsatisfactory. ☹

###### S1.1.8 RNAsamba

**RNAsamba** predicts the protein coding potential of RNA sequences using a neural network that incorporates variables from both RNA sequence and ORFs [13]. RNAsamba classifies input RNA sequences using canonical sequence features (ORF length, nucleotide k-mer frequencies, and amino acid frequencies) and a natural language processing neural network that uses the IGLOO architecture to identify informative motifs within the sequences without human input.

The training and test datasets employed are derived from multiple previously published datasets, these were CPC2 data (CCDS/RefSeq for mRNAs and GENCODE annotations of lncRNAs for a negative set from *H. sapiens*, *M. musculus*, *D. rerio*, *D. melanogaster*, *C. elegans* and *A. thaliana*). A FEELnc dataset from GENCODE (v24) *H. sapiens* transcripts with biotypes 'protein_coding' or 'lincRNA' & 'antisense' for the negative set. A "mRNN" dataset derived from GENCODE (v25) contained a "regular" set, and a challenging set ($\leq$ 50 codon short ORFS, and $\geq$ 50 codon untranslated ORFs in a lncRNA set). The total number of sequences combining CPC2, FEELnc and mRNN datasets is 12,142 coding and 18,019 non-coding sequences. It was mentioned that transcript associated with loci present in test data were excluded from training data, but sequence similarity between these sets did not appear to have been controlled, other than for the truncated ORF dataset where MMseqs2 was used to remove sequences from the training set with $> 90\%$ identity and coverage with the test set. Summary statistics such as length and C+G content were not reported for the positive and negative datasets. The reported average of performance over 5 species of RNAsamba a pre-trained model was 0.9915 sensitivity, 0.8891 PPV and 0.8639 MCC. We further note that only the forward strand (frames 1, 2 and 3) are evaluated by this tool, and the coordinates of predicted ORFs are not provided in the output.

```
$ rnasamba classify 'outputFile' 'input.fa'         'partial_length_weights.hdf5'
$ rnasamba classify 'outputFile' 'input-REVCOMP.fa' 'partial_length_weights.hdf5'
```

Installation documentation is available on their github code and io pages. Instructions are concise and allow for two routes to install, via pip or conda. Install options allow for installing a GPU version of the tool. Installation is excellent. ☺

Usage documentation contains instructions along with an example. Options were few but described. Error messages were useful. User experience is excellent. ☺

Output consists of a coding score and classification. Options allow to provide a protein fasta file output, however the output file does not contain frame information or any start and stop locations. Collating output data is simple as the output files contain little information and data is whitespace separated. Output is acceptable. 😐

###### S1.1.9 tcode

**tcode** implements the Fickett TESTCODE statistic and is included in the EMBOSS suite of bioinformatic tools [14, 15]. This is a composition-based statistic that considers the frequency of each nucleotide in turn in each codon position. The statistical parameters for the model were estimated and evaluated on 321 coding and 249 non-coding viral, bacterial, *Homo sapiens* and *Saccharomyces cerevisiae* sequences that were available in 1982.

The command-line parameters used for this evaluation were:

```
$ tcode  -window 200 -outfile outputFile input.fa
```

Tcode is part of the EMBOSS suite. The documentation for installing EMBOSS is very detailed and complex and does not provide a simple install. We decided to install tcode using bioconda. Installation is acceptable. 😐

Usage documentation is extensive and covers general and specific use. Multiple options are clearly documented and examples are given. Error messages are useful. User experience is excellent. ☺

Documentation on output is extensive. The output is verbose, but does make it difficult to collate multiple runs. It provides sliding window scores and does not provide start and stop sites for coding regions precisely. Output is acceptable. 😐

###### S1.1.10 Alignment-based tools:

###### S1.1.11 PhyloCSF

**PhyloCSF** "Phylogenetic Codon Substitution Frequencies" is a maximum likelihood estimate for a given input alignment and phylogenetic tree using a 64x64 codon rate matrix trained on either coding or non-coding models [16]. Alignments are scored against both models using a log-likelihood ratio ($\log\left(\frac{p_c}{p_n}\right)$). PhyloCSF was recently reimplemented in C++, this new implementation "PhyloCSF++" is expected to be easier to install, and faster; However, it lacks features like an all-frame scanning option [17]. While PhyloCSF has modes that allow training on the specific dataset being studied, this option conflicts with our inclusion criteria. Therefore, we have used the generic "Omega Test" mode that requires a phylogenetic tree and not a species-specific empirical codon models that require the estimation of thousands of parameters. This mode is closely related to the widely used $\frac{d_N}{d_S}$ likelihood test. Based on the documentation, it is unclear how this model was parameterised, and whether positive and negative datasets were used in the development. While PhyloCSF performance measures were not reported directly in the manuscript, we can estimate these from ROC curve for "12-species alignments" and the "full dataset" shown in Figure 2A of the manuscript [16], the specificity is approximately 0.98 and the sensitivity is approximately 0.925. We further note that the coordinates of predicted ORFs are not provided in the output.

The command-line parameters used for this evaluation were:

```
$ PhyloCSF -f6 --removeRefGaps --strategy=omega modelFile input-alignment.fa
```

Installation instructions are substantial, but very complex. It requires installing a package manager for OCaml, then installation of multiple dependencies. It requires the user to compile PhyloCSF. We could not get OCaml to install correctly, which means we could not build PhyloCSF and install it using the provided instructions. We switched to using bioconda. Installation is unsatisfactory. ☹

Usage documentation is extensive, detailing user options, alignment preparation and provides multiple examples. A phylogenetic tree is required to use the "Omega" mode, which is a generalised coding score model. The tree and alignments must have exact naming and be formatted perfectly, and error messages did not help in identifying formatting as an issue. User experience is acceptable. 😐

Documentation on output is also extensive, detailing output metrics. The output contains the highest scoring region and corresponding strand, however it does not report start and stop coordinates. The strand start and alignment length are also provided. Extracting information from the output was easy due to simple tab-delimitted formatting. Output is acceptable. 😐

###### S1.1.12 RNAcode

**RNAcode** [18]is a comparative analysis tool that identifies "evolutionary signatures" of protein coding sequences. A dynamic programming algorithm is used to find locally maximal scoring regions in an alignment with a reference sequence. The score aggregates information from nucleotide and amino acid substitutions as well as frame-preserving gaps. The algorithm is tolerant of alignment and sequencing error so frame-shift due to errors are not over-penalised. The statistical significance of high scoring pairs is evaluated by simulating neutral alignments with similar evolutionary constraints. While machine-learning was not used to train this method, it was evaluated on enterobacterial, archaeal, insect, nematode and vertebrate coding and non-coding regions. The self-reported performance metrics for RNAcode were based on an automated annotation of Drosophila genome alignments and Flybase annotations were 0.88 sensitivity and 0.93 specificity.

The command-line parameters used for this evaluation were:

```
$ RNAcode -s -t input.aln -o outputFile
```

Documentation is extensive and provides options for root or non-root install as well as detailing multiple options when installing. Unfortunately we could not get it to install due to compile errors and switched to using bioconda for installation. Installation is unsatisfactory. ☹

Usage documentation is extensive, detailing user options, alignment preparation and provides multiple examples. Options were highly useful, as were error messages. User experience is excellent. ☺

Documentation on output is extensive and provides options on how the data is formatted. This made it very easy to collate multiple output files. Start and stop sites of the coding region were provided as well as strand. This was also one of the few tools to provide a p-value for the scores given. Output is excellent. ☺

#### S1.2 Relative accuracy and runtime figures

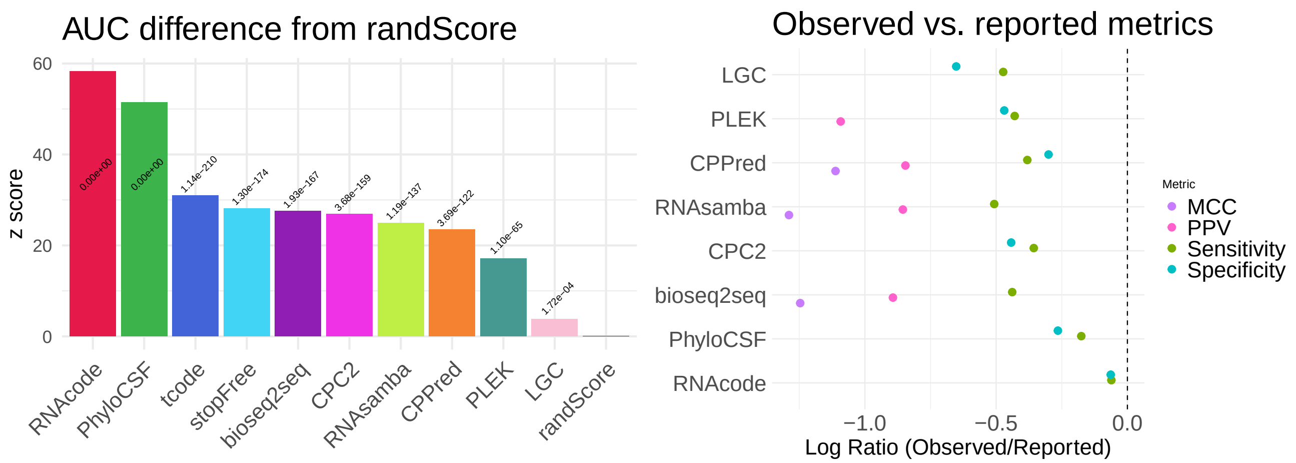

**Figure S1. Accuracy relative to a random number or self-reporting. A:** $z$ score differences in the area under the ROC curve for each tool compared to a random number generator, with corresponding p-values derived from bootstrap analysis. **B:** Log ratios comparing self-reported to independently measured tool accuracy, where values near zero indicate high agreement and more negative values indicate greater discrepancies.

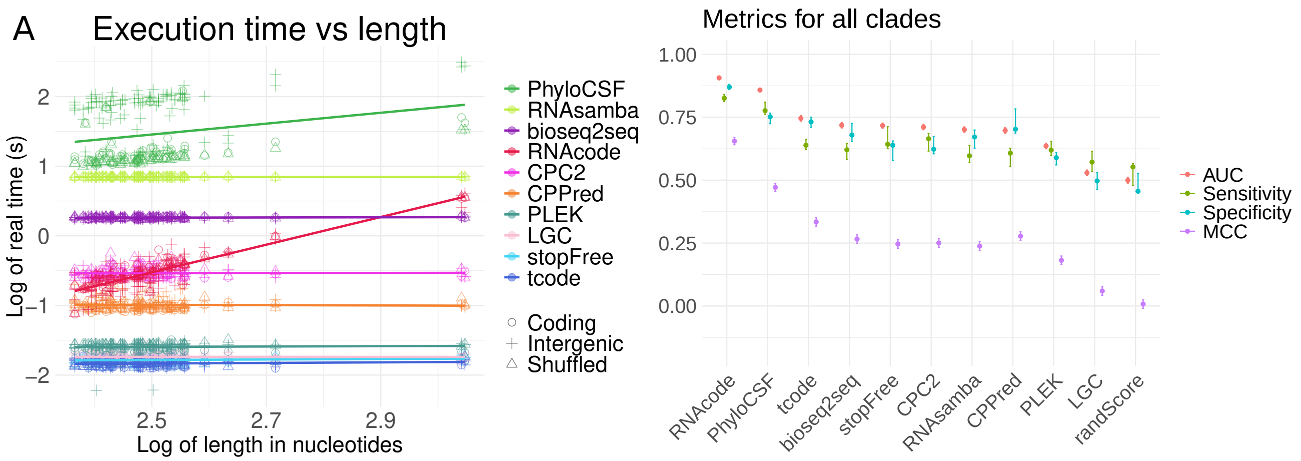

**Figure S2. Compute time and average accuracy. A:** Time vs length scatter plot. The regression lines are shown for each tool. The initialisation times for each tool is given by the $y$ intercept, the slope of the line shows the dependency of each tool on sequence length. **B:** Accuracy metrics of combined clades with 95% confidence intervals estimated by bootstrapping the results with 'pROC' (AUC, sensitivity, specificity) and 'boot' R functions. In each case 5,000 replicates with the same number of samples as the original datasets.

#### S1.3 Clade-specific result figures

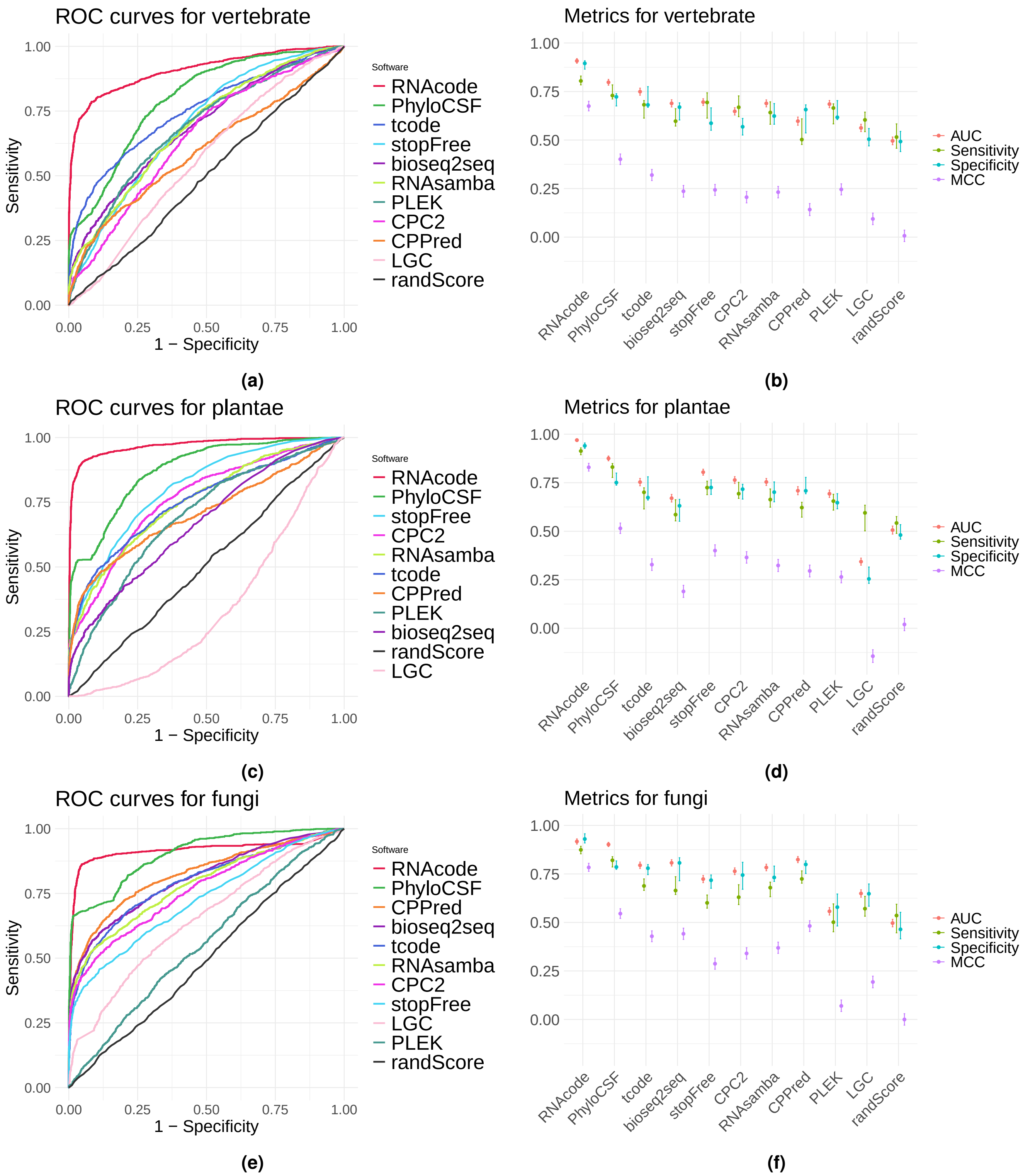

**Figure S3. Clade specific accuracy.** Accuracy comparisons for the vetebrate (**a**&**b**), plant (**c**&**d**) and fungi (**e**&**f**) datasets. ROC curves are shown in the left column (**a**,**c**&**e**), and point estimates of AUC, sensitivity and MCC metrics are shown in the right column (**b**,**d**&**f**).The 95% confidence intervals are indicated with whiskers on each point estimate. These were estimated by bootstrapping the results with 'pROC' (AUC, sensitivity, specificity) and 'boot' R functions. In each case 5,000 replicates with the same number of samples as the original datasets.

#### S1.4 Score distributions figures

In order to compare score distributions $\tilde{s}$ for each tool we use a $z$ score normalised for each tool. Within each clade a mean ($\bar{x}$) and standard deviation ($\sigma$) is computed from tool scores ($x$) for pooled negative controls of intergenic and shuffled sequences or alignments. To better cater for extreme distributions, the scores for bioseq2seq, CPC2 and CPPred were transformed with a $\log_{10}$ function.

$$z = \frac{x - \bar{x}}{\sigma}$$

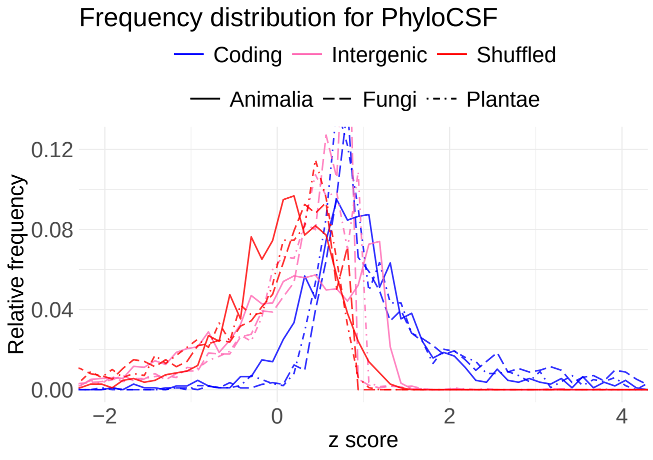

**(a)**

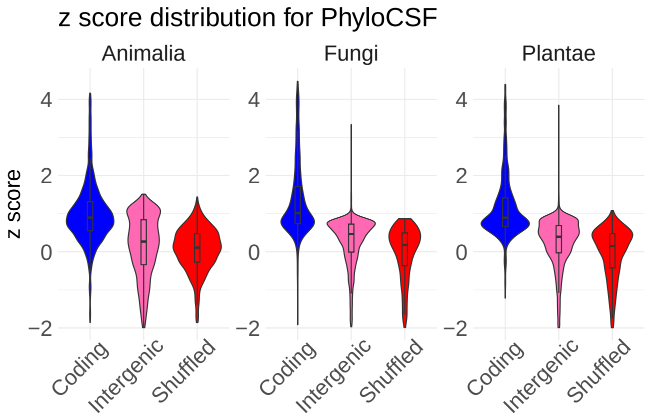

**(b)**

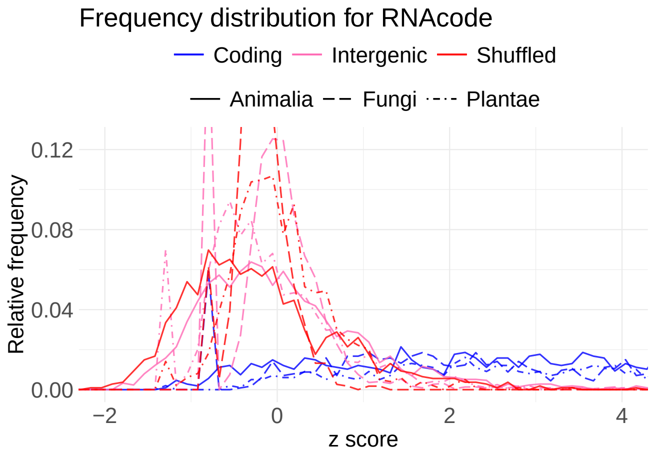

**(c)**

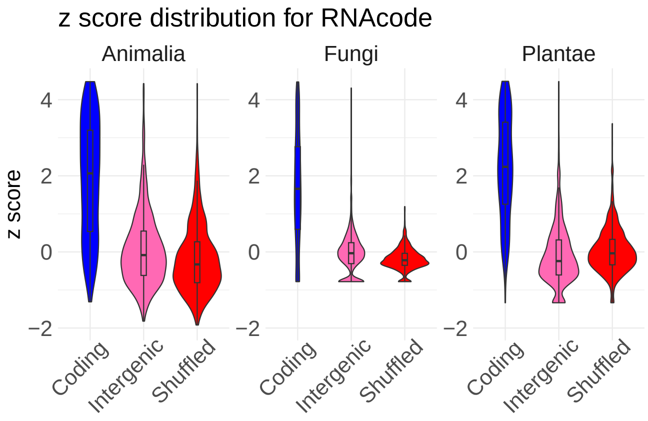

**(d)**

**Figure S4.** Score distributions for alignment-based tools PhyloCSF (first row panels **a** & **b**) and RNAcode (second row panels **c** & **d**). The $z$ scores for coding, intergenic, and shuffled alignments have been calculated for each test clade. The distributions are depicted as histogram line graphs (left panels **a** & **c**) and violin plots (right panels **b** & **d**) for each clade and data type.

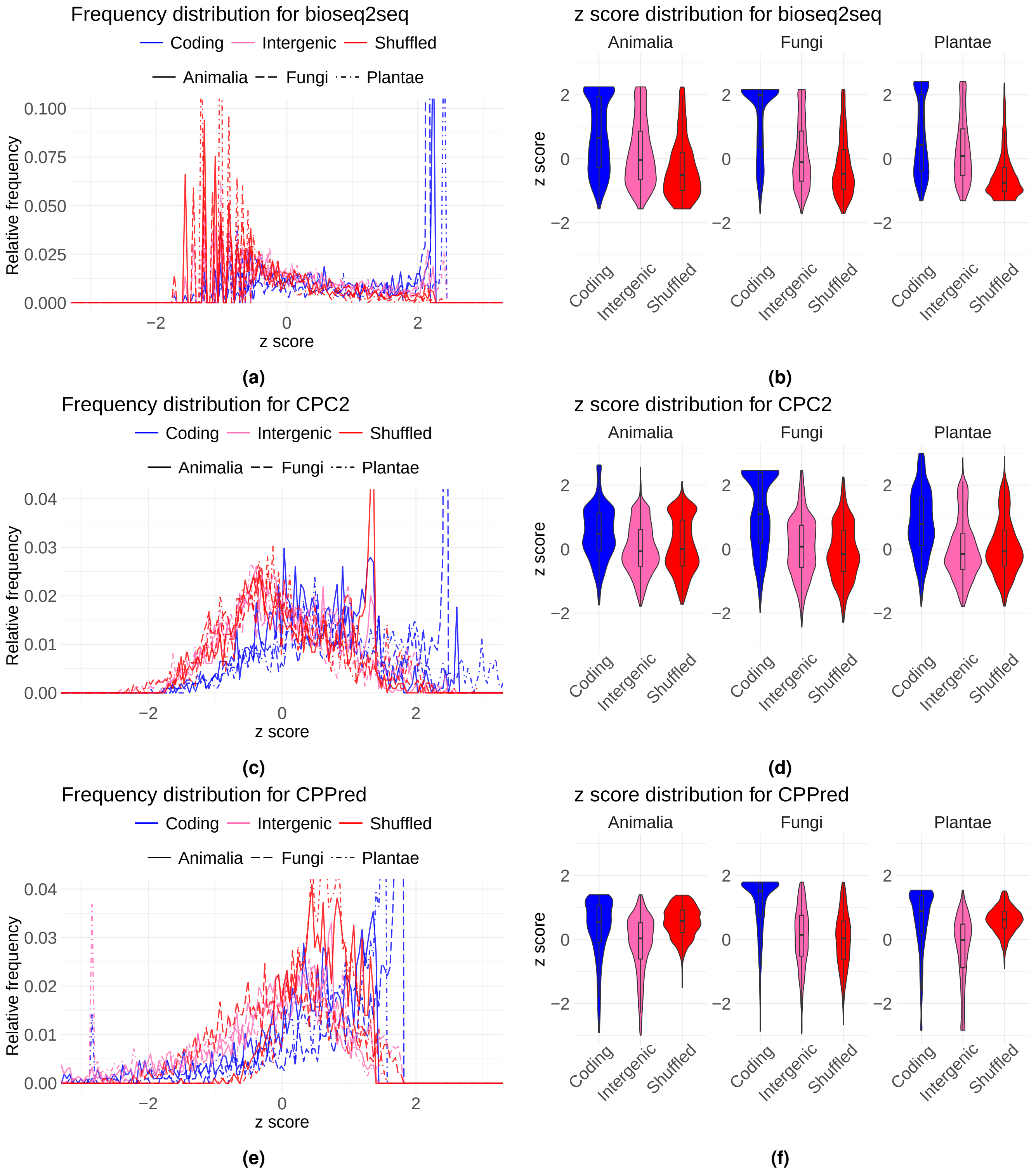

**Figure S5.** Score distributions for sequence-based tools bioseq2seq (first row panels **a** & **b**), CPC2 (second row panels **c** & **d**) and CPPred (third row panels **e** & **f**). The $z$ scores for coding, intergenic, and shuffled sequences have been calculated for each test clade. The distributions are depicted as histogram line graphs (left panels **a**, **c** & **e**) and violin plots (right panels **b**, **d** & **f**) for each clade and data type.

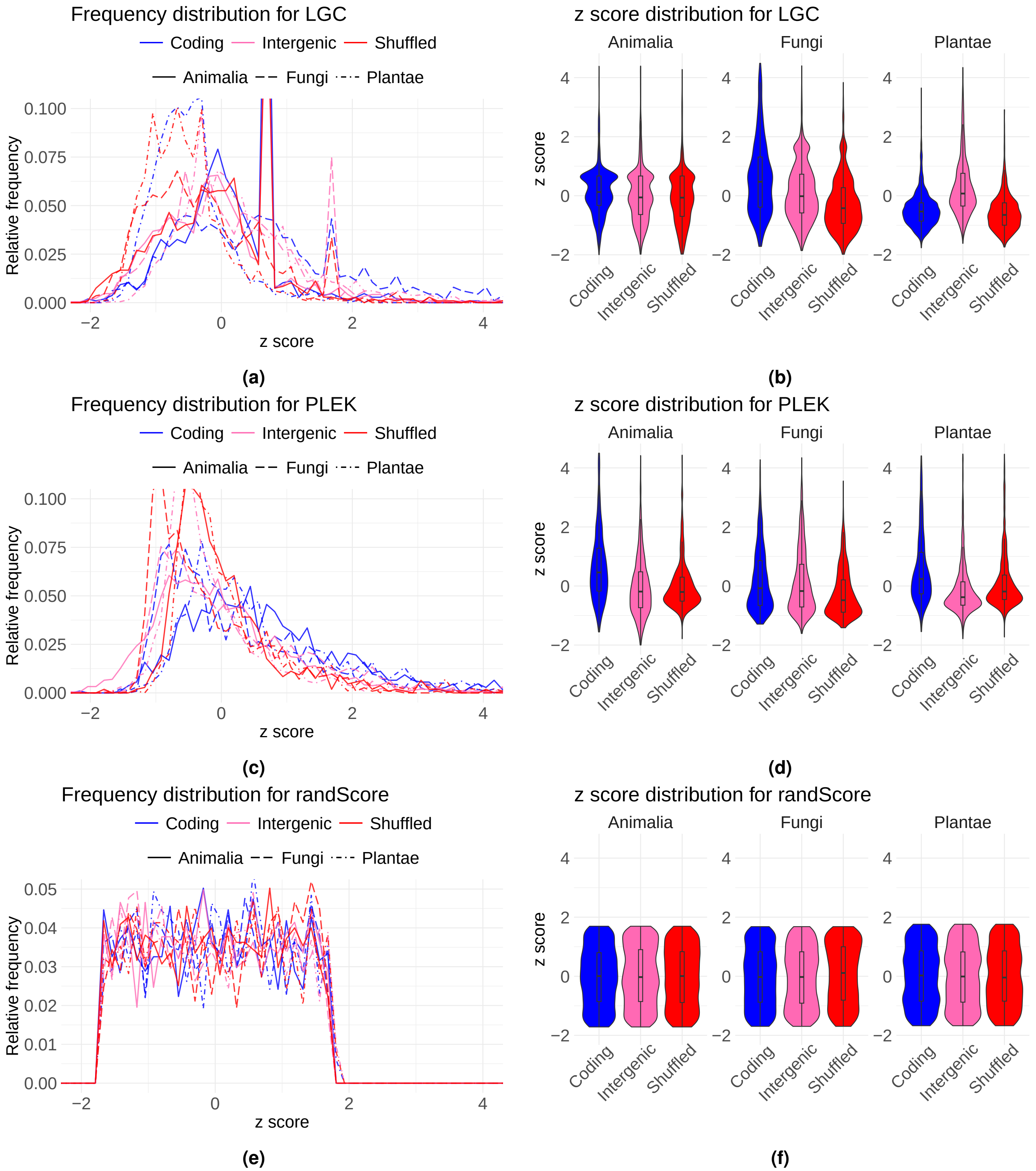

**Figure S6.** Score distributions for sequence-based tools LGC (first row panels **a** & **b**), PLEK (second row panels **c** & **d**) and randScore (third row panels **e** & **f**). The $z$ scores for coding, intergenic, and shuffled sequences have been calculated for each test clade. The distributions are depicted as histogram line graphs (left panels **a**, **c** & **e**) and violin plots (right panels **b**, **d** & **f**) for each clade and data type.

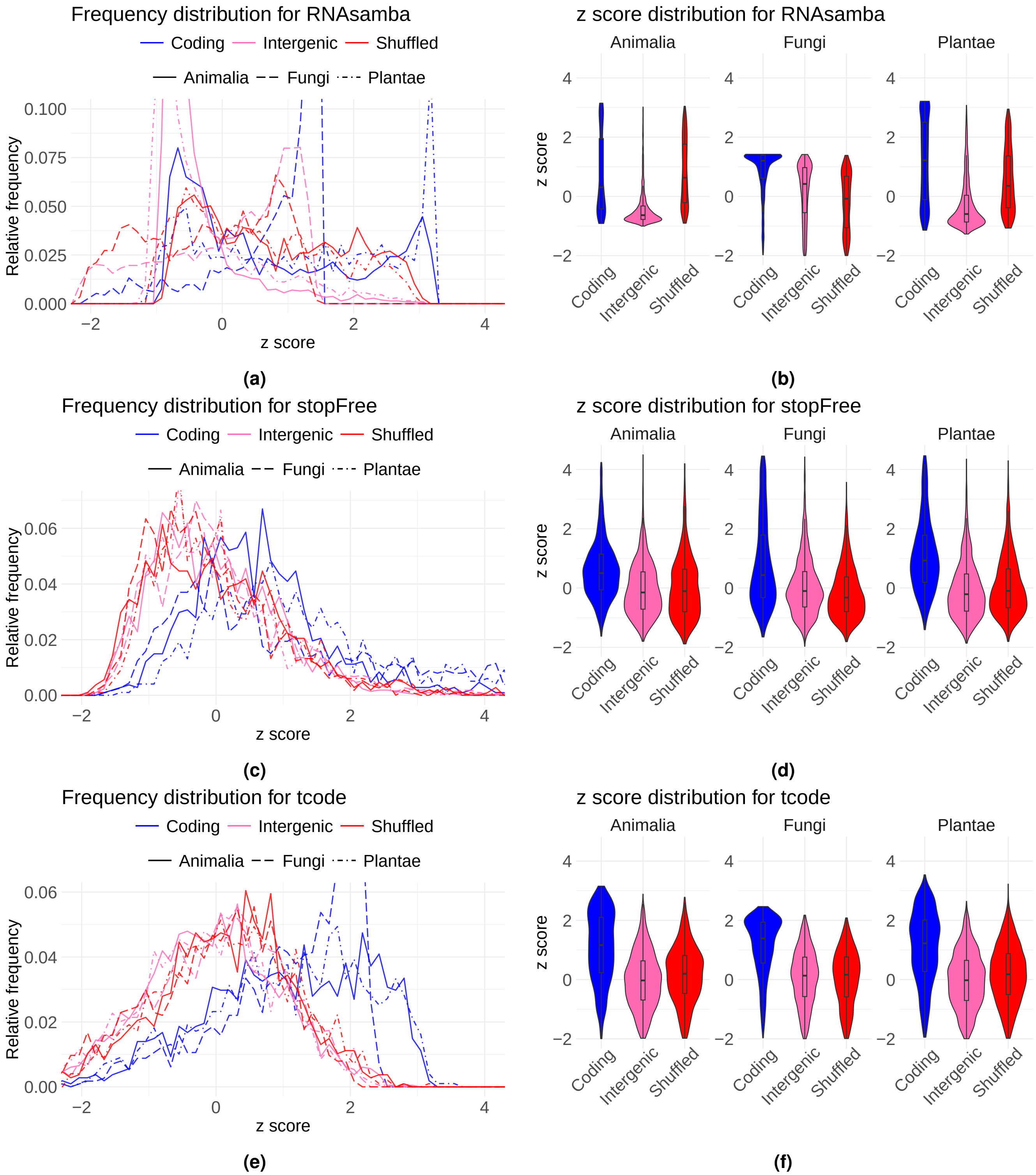

**Figure S7.** Score distributions for sequence-based tools RNAsamba (first row panels **a** & **b**), stopFree (second row panels **c** & **d**) and tcode (third row panels **e** & **f**). The $z$ scores for coding, intergenic, and shuffled sequences have been calculated for each test clade. The distributions are depicted as histogram line graphs (left panels **a**, **c**, & **e**) and violin plots (right panels **b**, **d**, &**f**) for each clade and data type.

### 2. Benchmarking on Short ORFs

#### S2.1 sORF Benchmarking

Accurate annotation of Short Open Reading Frames (sORFs) ($\leq$ 300 nts, $\leq$ 100 amino-acids) is challenging due to the limited length [19]. This can lead to their misclassification by standard annotation pipelines, despite their recognition for diverse biological roles [20, 21]. To assess tool performance on these challenging sequences, we used a curated set of approximately 40 sORFs for each of the clade used in the main study.

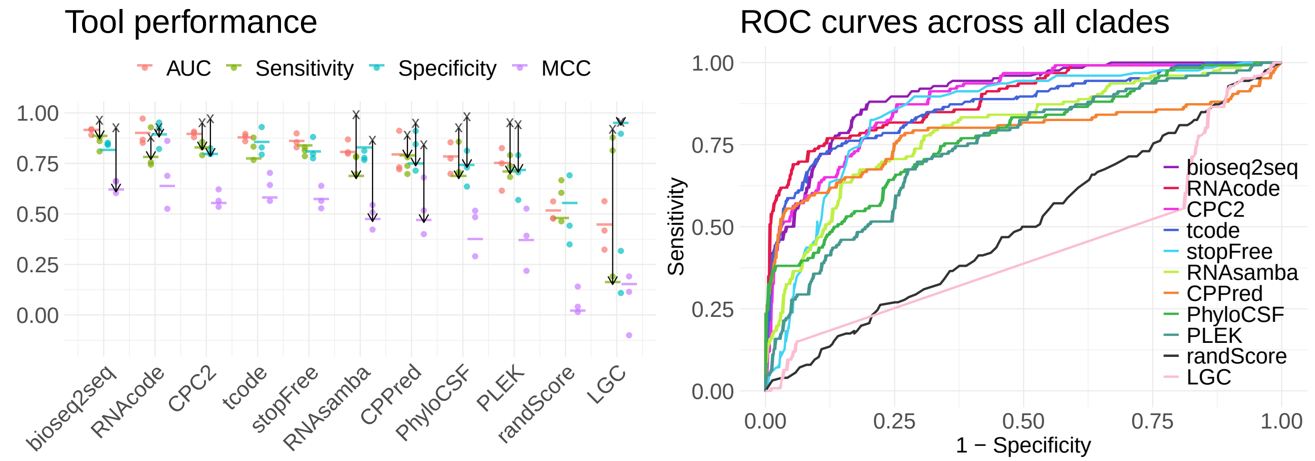

**Figure S8. Coding potential tools, accuracy on sORFs A:** A jitter plot showing the mean AUC, Sensitivity, Specificity and MCC values for each software tool and clade when tested on sORFs. Tools are ordered by the median AUC (mean performance values are indicated with a tick mark for each metric and tool). The black crosses and arrows indicate the accuracy metrics reported by the authors of each tool on their own test datasets. **B:** ROC curves showing the ability of each tool to discriminate between positive coding sequences and negative unannotated, length matched genome regions, and shuffled controls using the sORF dataset.

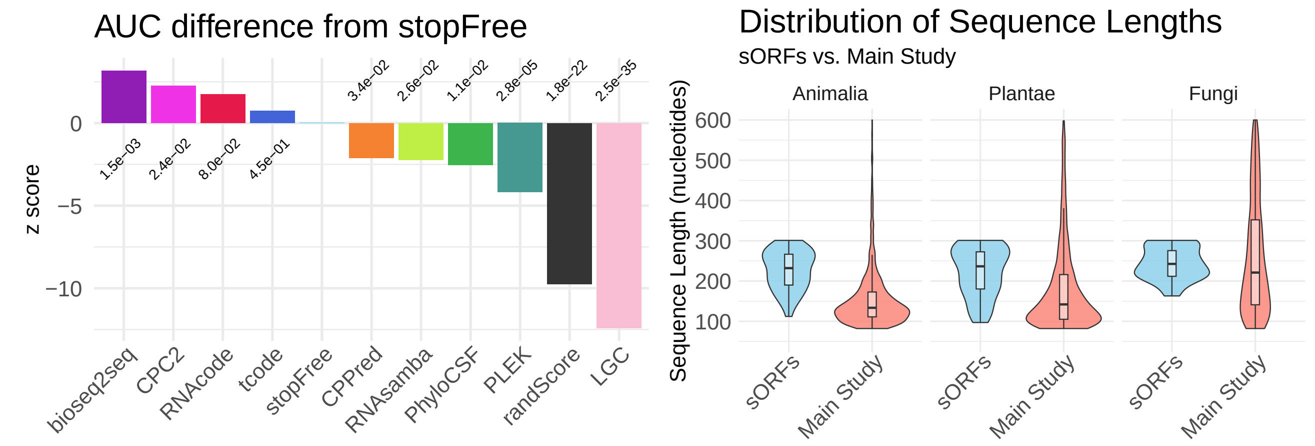

**Figure S9. Accuracy relative to stopFree A:** z score differences in the area under the ROC curve from our naive stopFree tool, with notated p-values when run on the sORF dataset. **B:** Distribution of sequence lengths controls used in the main study and sORF benchmarks.

[9] Bi-Qing Li, Kai-Yan Feng, Lei Chen, Tao Huang, and Yu-Dong Cai. Prediction of protein-protein interaction sites by random forest algorithm with mrmr and ifs. *PLoS ONE*, 2012.
